# DystoGen Compendium: A comprehensive resource of ACMG annotated movement disorder associated genetic variants

**DOI:** 10.1101/2023.10.31.564874

**Authors:** Bhaskar Jyoti Saikia, Utkarsh Gaharwar, Mukta Poojary, Aditi Mhaske, Sangita Paul, Mukesh Kumar, Srishti Sharma, Kavita Pandhare, Shivanshi Rastogi, Medhavi Karal, Mercy Rophina, Vinod Scaria, Binukumar BK

## Abstract

**Purpose:** In recent years, the advent of high throughput sequencing techniques has led to the identification of a number of genetic variants across different genes that are associated with movement disorders. However, the under-appreciation of the variant spectrum in movement disorders and the lack of consolidated and systematic evidence-based annotation of these variants has long undermined the true potential of genomic approaches to expedite precision medicine.

**Methods:** We manually curated the genetic variants from a panel of 118 genes that have been associated with monogenic causes of movement disorders and systematically annotated them according to ACMG & AMP (American College of Medical Genetics and the Association of Molecular Pathologists) guidelines.

**Results:** Data integration after systematic classification of variants according to ACMG & AMP guidelines showed 5118 pathogenic/likely pathogenic variants accounting for 18.03% of the total unique variants being annotated. This data and annotations are available in a comprehensive online compendium DystoGen.

**Conclusion:** To the best of our knowledge, this is the most comprehensive compendium of genetic variants in movement disorders annotated as per the ACMG & AMP guidelines for pathogenicity. The compendium indexes 28377 variants along with a wide array of information including the geographical origin of the variant, global distribution, and population allele frequency. The resource has been made available in the URL https://clingen.igib.res.in/dystogen/.

## Introduction

Movement disorders are a class of heterogeneous group of neurological complications that are diverse in terms of genetics, pathology and clinical characteristics, associated with abnormal bodily movements. These disorders can cause a wide range of symptoms, including tremors, stiffness, slowness of movement, and problems with balance and coordination. Movement disorders can be caused by a variety of factors, including genetics, brain injury, and exposure to toxins. Despite this variation, there is a great deal of overlap across the many types of movement disorders because they all share the same flaws, such as problems with movement planning, control, or execution in common (1). The motions may be voluntary or involuntary and have been linked to a variety of neurological conditions, but is not limited to Dystonia, Amyotrophic Lateral Sclerosis, Ataxia, Parkinson’s disease, tremors, dyskinesia, and others (2). Clinical manifestations of movement disorders can be complicated, inconsistent, and occasionally odd. It can therefore be challenging to make the right diagnosis, even in the hands of trained movement disorder specialists. But precise recognition based on clinical insight is crucial for a number of reasons.

For the subsequent diagnostic process to be successful, the type of movement disorder must first be correctly classified. The majority of diseases lack a particular biochemical signature that may clearly identify the underlying illness, with subsequent overlap in symptoms with other disease types (3,4). Recent genetic studies have elucidated the involvement of multiple genes in causing different types of movement disorders, but the genetic heterogeneity and frequent clinical overlapping of phenotypic manifestations of various movement disorders renders difficulty in clinical settings for efficient diagnosis and devising treatment regimens (5)

Over the past few decades, significant developments in clinical diagnostic testing have been made, especially in the understanding of the genetic variants associated with genetic causes of various movement disorders. Consequently, clinical genetic testing in recent years has shown a promising opportunity for early and precise diagnosis of different heritable forms of movement disorder (6). Contrary to the single-gene sequencing approach for genotype-phenotype correlation studies and clinical genetic testing of patients, strategies such as targeted gene panel analysis and whole-exome sequencing (WES) approach have proven to be satisfyingly useful in diagnosing complex neurological disorders in patients, in which the underlying genetic etiology remains unclear (7–10). *in-vivo* and *in-vitro* studies have also aided in understanding not only the cause and effect of associated genetic variants but have also helped in understanding the underlying pathological pathways and the direct or indirect association of certain genes in the disease progression (11–13). Despite these achievements, the lack of a unified comparative resource and recommendations for systematic annotation of genetic variants have long undermined the true potential of such robust diagnostic tools. Recently, the American College of Medical Genetics and the Association of Molecular Pathologists (ACMG & AMP) have jointly put forward a guideline for the systematic annotation of genetic variants and recommendations for the classification of variant’s pathogenicity based on available evidence (14).

In the present study, we re-evaluated the pathogenicity of a comprehensive set of genetic variants curated from literature as well as databases in the public domain according to ACMG & AMP guidelines. We broadly curated a panel of 118 genes associated with various types of movement disorders, with a total of 28377 variants being reported globally, based on available evidence from the scientific literature, commercial gene panels, and reputable databases such as MDSGene, ClinVar, LOVD (Leiden Open Variation Database), to comprehensively encompass a broad range of movement disorders. The genes were categorized into various categories as listed in **Supplementary Data 1** based on their involvement in a particular disease with a specific disease manifestation. We also attempted to understand the allele frequencies of Pathogenic and Likely pathogenic variants in global population datasets. This dataset is also available as a searchable resource - Dystogen is available in the URL https://clingen.igib.res.in/dystogen/.

## Materials and Methods

### Data Sources

Literature Sources: The published literature was queried for relevant literature on the genetic causes of movement disorders. Literature resources - Pubmed Central (PMC) and Pubmed, were queried using keywords that include the terms variant, mutation, polymorphism, gene name, and associated disease. Only English language literature was considered for the curation.

Database Sources: A number of open-source datasets and databases were searched for genes and variants associated with genetic dystonia. This included the Leiden Open Variation Database (LOVD), ClinVar, and MDSGene Database. The source of variants information has been listed in a tabular form.

### Annotation of variants

While generating a database, the most important step is proper validation of the curated data. To maintain the uniformity and unambiguousness of the data, a routine validation check is inevitable. In the case of genetic variants, data validation was performed with the help of two web-based tools, Mutalyzer (https://mutalyzer.nl/) and Variant Validator (https://variantvalidator.org/).

### Mutalyzer

It is a web-based interface that is widely used for validating, constructing, converting sequence variant descriptions and aids in extracting missing shreds of information for the genetic variants. The tool was specifically used to convert a chromosomal position of a variant description into a position relative to a RefSeq transcript reference sequence of a local database (NCBI) or vice-versa with the help of the position converter feature (15).

### Variant Validator

The variant validator is a web-based interface that strictly complies with HGVS (Human Genome Variation Society) Sequence Variant Nomenclature for variant description, validation, and reporting syntax errors if made by the end-user. The tool was used to validate the variants collected from different sources and were checked for errors if reported followed by correction with the help of the UCSC browser. Upon validation, HGVS-compliant variant descriptions were reported with all available reference sequences, variant description in gene level (RefSeqGene), and chromosomal position in both HGVS and VCF format (16).

### Annotation of variants using ANNOVAR

An in-silico computational tool, ANNOVAR (ANNOtate VARiation) was used to functionally annotate the validated genetic sequence variants in VCF (Variant Calling Format) format, containing five basic fields: chromosome, start and end position of the nucleotide, reference, and altered nucleotides. The functional annotation consists of three subtypes: Gene-based annotation, Region-based annotation, and Filter-based annotation. These annotations were systematically used to identify if the input variants are exonic, intronic, intergenic, splice site, 3’/5’ UTR, upstream/downstream of genes or causing protein-coding change due to mutation and also to obtain variants from databases such as Esp6500, ExAc, and 1000 Genome Project, obtaining allelic frequencies and in-silico prediction scores generated by SIFT, PolyPhen, and CADD for determining the deleteriousness of the sequence variant (17).

### Data Curation

The collected datasets containing relative parameters were consolidated into an Excel spreadsheet to include 1) Genome Reference Consortium Human build 37 (GRCh37/hg19, 2) Reference, 3) HGVS_NM, 4) Chromosome, 5) Amino Acid Change, 6) Reference Gene, 7) Population, 8) Disease, 9) OMIM, 10) dbSNP, 11) HGVS_NP, 12) Reference base, 13) Altered base, 14) Start position, 15) End position, 16) Homozygous/Heterozygous, 17) Genetic Origin, 18) Technique 19) Ethnicity, 20) Geographical Origin, 21) HGVS_NC, 22) HGVS_NG, 23) Inheritance, 24) ClinVar ID, 25) Mutation, 26) ClinVar_SIG, 27) MutationEffect.

### Classification of the pathogenicity of variants according to the ACMG-AMP guidelines

The variants were systematically analyzed and classified according to the guidelines developed by ACMG & AMP (14). This guideline reports a standard for variant classification as Benign, Likely benign, Pathogenic, Likely pathogenic, and variants of uncertain significance (VUS) for identification of variants. This classification is based on 28 criteria followed for sequence variants that include population allele frequency, computational data, segregation data, variant type, and literature screening for functional studies data. The following attributes were used for variant classifications: **PVS1-** Nonsense, Frameshift or Splicing variant in a gene in which the consequences of its loss of function (LOF) for a disease phenotype is well established. **PS1-** Same amino acid change, irrespective of nucleotide change, previously established as a pathogenic variant. **PS2-** De Novo mutation (confirmed paternity and maternity) in a patient. **PS3-** Substantial evidence from functional studies in in-vitro and in-vivo model systems shows damaging effects of the variant under consideration. **PP1** (Segregation evidence)- Co-segregation of the variant with the disease in multiple affected relatives that is positively known to cause the disease. Variant segregating in a single, two and more than two generations of family were given PP1, PP1- Moderate (PP1-M) and PP1-Strong (PP1-S) respectively. **PM1**- Variant present in functionally important protein domain. Based on the allelic frequency of the variants from the 3 databases that include Exome Sequencing Project 6500 (Esp6500), 1000 Genomes Project, and Exome Aggregation Consortium (ExAC), variants were classified into **BA1-** Allele frequency greater than or equal to 0.05, **BS1**- 0.005-0.05. **BS2**- 0.0005-0.005 and **PM2**- less than or equal to 0.0005. **PM3 (Compound Heterozygous)**- Detected in trans with a previously well-established pathogenic variant for recessive disorders. **PM4**- Change in the protein length as a consequence of indels (inframe insertions and deletions) that is not present in the repeat regions. **PM5**- Novel point mutation at an amino acid residue in which another missense change has been established as a pathogenic variant previously (same position but different amino acid). **PM6**- De novo variant and identity of parents is assumed. **PP2**- Missense variant in a gene known to have missense variants as a frequent cause of disease. Based on the 3 computational tools SIFT, Polyphen, and CADD score, the 2 attributes assigned were **PP3**- 2/3 tools predict pathogenicity (**SIFT Pred = Damaging or SIFT score < 0.05, Polyphen2_HVAR_Pred = Damaging or Polyphen2_HVAR_ score > 0.446, CADD score > 15**) and **BP4**- 2/3 predict benign (**SIFT Pred = Benign/tolerant or SIFT Score > 0.05, Polyphen2_HVAR_Pred = Benign/tolerant and Polyphen2_HVAR_score ≤ 0.446, CADD Score ≤ 15**). **PP4-** Clinical sensitivity of testing is high, well-defined syndrome with little overlap with other clinical presentations, genes do not harbor a lot of benign variants, family history is consistent with the mode of inheritance of disease observed **BS3**- Substantial evidence from functional studies showing a tolerable/benign effect. **BS4**- Non-segregation of the variants in affected members of a family with the disease. **BP1**- Nonsense variant in a gene with a low rate of pathogenic nonsense variation.**BP2**- Observed in cis with a pathogenic variant irrespective of its inheritance pattern or in trans with a dominant pathogenic variant. **BP3**- Inframe indel variants in the repeat region of a gene. **BP4**- In-silico computational tools predict the benign effect of a variant. **BP6**- Well-known source reporting the variant to be benign but without well-established functional studies. **BP7 -** Synonymous variant not known/predicted to have any effect on splicing.

### Comparison of variant frequencies with global populations

Of the total variants, only those variants which were categorized as pathogenic or likely pathogenic by the ACMG-AMP guidelines were considered for further analysis. P/LP variants which were found in the Indian population were systematically filtered by comparing with the Indigenomes resource (Jain et al., 2021). Minor allele frequencies of these variants were retrieved for major global populations including Africans, Americans, Europeans (Finnish and non-Finnish origin), Middle Easterns, Amish, East Asians and South Asians from the gnomAD data resource and were compared (Karczewski et al., 2020). Statistical significance was estimated using Fisher’s exact test and Bonferroni multiple corrections (p-value ≤ 0.05).

### Data processing and statistical analysis

The initial level of data processing involved fetching all variants which were classified as pathogenic or likely pathogenic as per the ACMG guidelines. This encompassed a total of 5118 variants and corresponds to human genome 19 (GRCh37/hg19) assembly. The genetic variant coordinates were systematically lifted over to GRCh38/hg38 assembly using NCBI remap. Global population frequencies of the variants were retrieved from the gnomAD v2.1.1 dataset which included exome and whole genome data from Indian (Indigenome) as well as major global populations as mentioned in the previous section using bespoke scripts.

Global population frequencies were fetched, and statistical significance was estimated using Fisher’s exact test and Bonferroni multiple corrections *(p-value ≤ 0.05).* Minor allele frequencies of statistically significant variants among the Indian and global populations are shown in Figure 3.

### Database and web server

A web-based search tool was integrated with the DystoGen database to allow the users to easily explore the genes and variants. The database was converted into JavaScript Object Notation (JSON). All variants were stored into MongoDB v3.4.10 and the server was hosted using Apache HTTP server. The user-friendly web interface for querying the database was coded in HTML5, CSS3, Bootstrap version 4.0 (Material Design), and AngularJS. The back-end of the database was constructed using PHP 7.0 and MongoDB v3.4.10. MongoDB v3.4.10 was used to keep track of data processing and database through the web interface. The search query was optimized using indexing in MongoDB, with most search terms, including gene names, variant, and dbSNP ID populating the search bar. AngularJS and PHP 7.0 scripts were used to retrieve and bind the data from the database.

## Results

### Curation of genes and variants

In the present analysis, we systematically collected and curated a total of 34180 variants encompassing 118 genes associated with dystonia, ataxia, and other movement disorders, curated from different databases and published literature in the English language. Of the total 34180 variants, a total of 28377 variants were found to be unique and were taken up for evaluation of evidence as per the ACMG and AMP guidelines. The variants in the database were broadly classified as Substitution (24757) (87.24%), Insertion (1138) (4.13%), Deletion (2292) (8.04%) and Indels (190) (0.67%), shown in Figure 1A.

**Figure 1:**
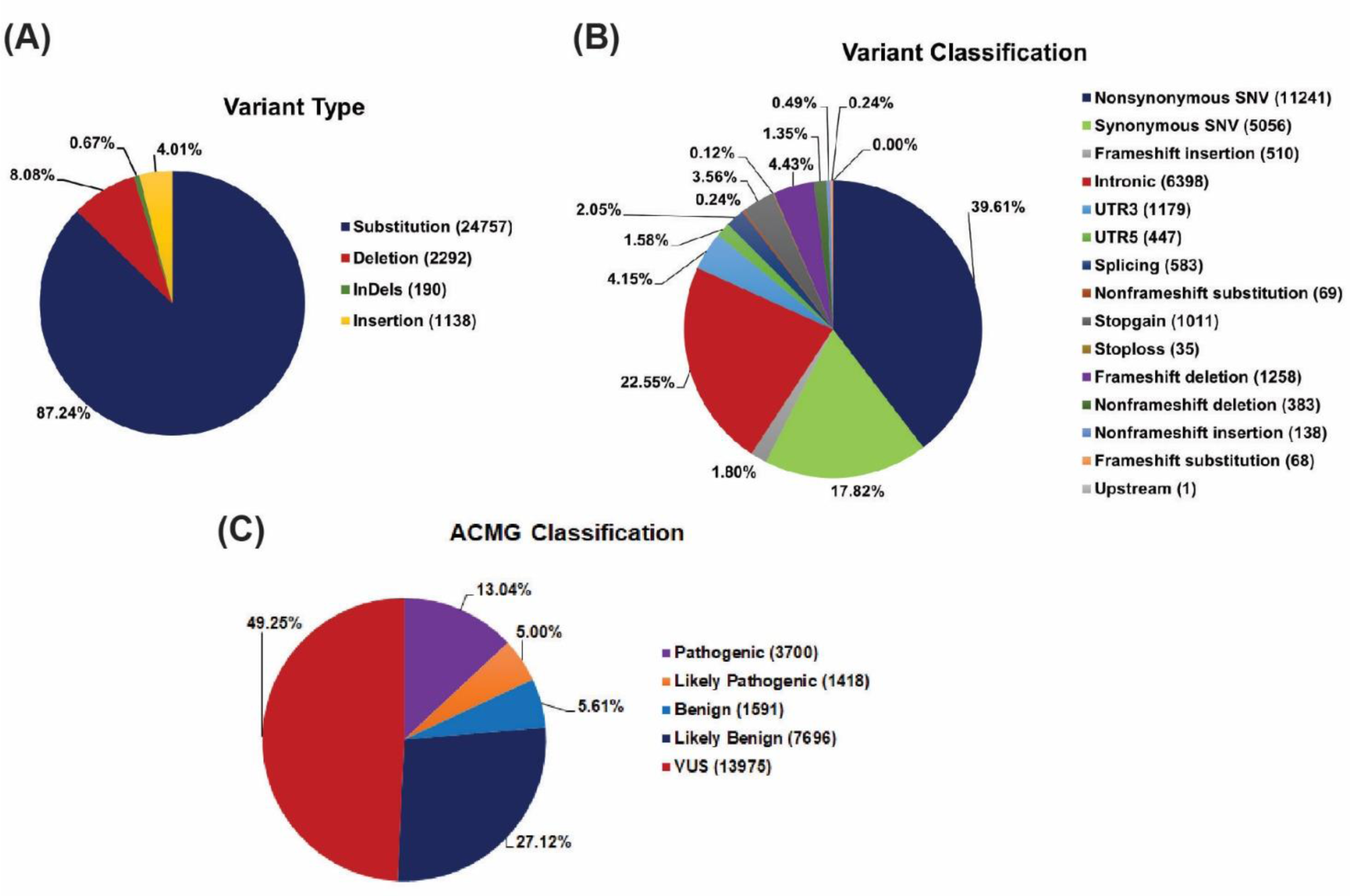
Classification of variants as (a) Variant Type, (b) Variant Classification, and (c) ACMG Classification.

The variants were further classified according to their functional implications, which have been designated under the category variant classification as shown in Figure 1B. Nevertheless, the validated and annotated variants were further classified according to the ACMG & AMP recommendations, of which 1415 were likely pathogenic (4.99%), 3700 were pathogenic (13.04%), 1591 were benign (5.61%), 7696 were likely benign (27.12%) and 13975 were variants with unknown significance (VUS) (49.25%). The ACMG & AMP classified genetic variants are depicted in Figure 1C.

### The genetic fabric of variations in different types of Dystonia

The present data encompasses 118 movement disorder associated genes. Broadly, as described earlier, dystonia is an amalgamation of various types of dystonia including myoclonus dystonia, focal dystonia, early-onset dystonia, late-onset dystonia, generalized dystonia, and other disease phenotypes, symptoms of which mostly overlaps with ataxia, parkinsonism, and other movement disorders, spanning across various genes. A stacked bar graph displaying genes belonging to different dystonia types and associated disease along with number of pathogenic/likely pathogenic variants associated with each gene has been shown in Figure 2. Here we report a total of 28377 unique variants, of which 1415 are Likely Pathogenic and 3700 are Pathogenic. Out of the total 5118 pathogenic and likely pathogenic variants, 1980 are present in the functional domains of the associated protein. Overall, the pathogenicity of dystonic variants at present is 18.02% according to the data being displayed. Further individual gene clusters with pathogenic/likely pathogenic variants of each gene have been tabulated in **Supplementary Data 2**.

**Figure 2:**
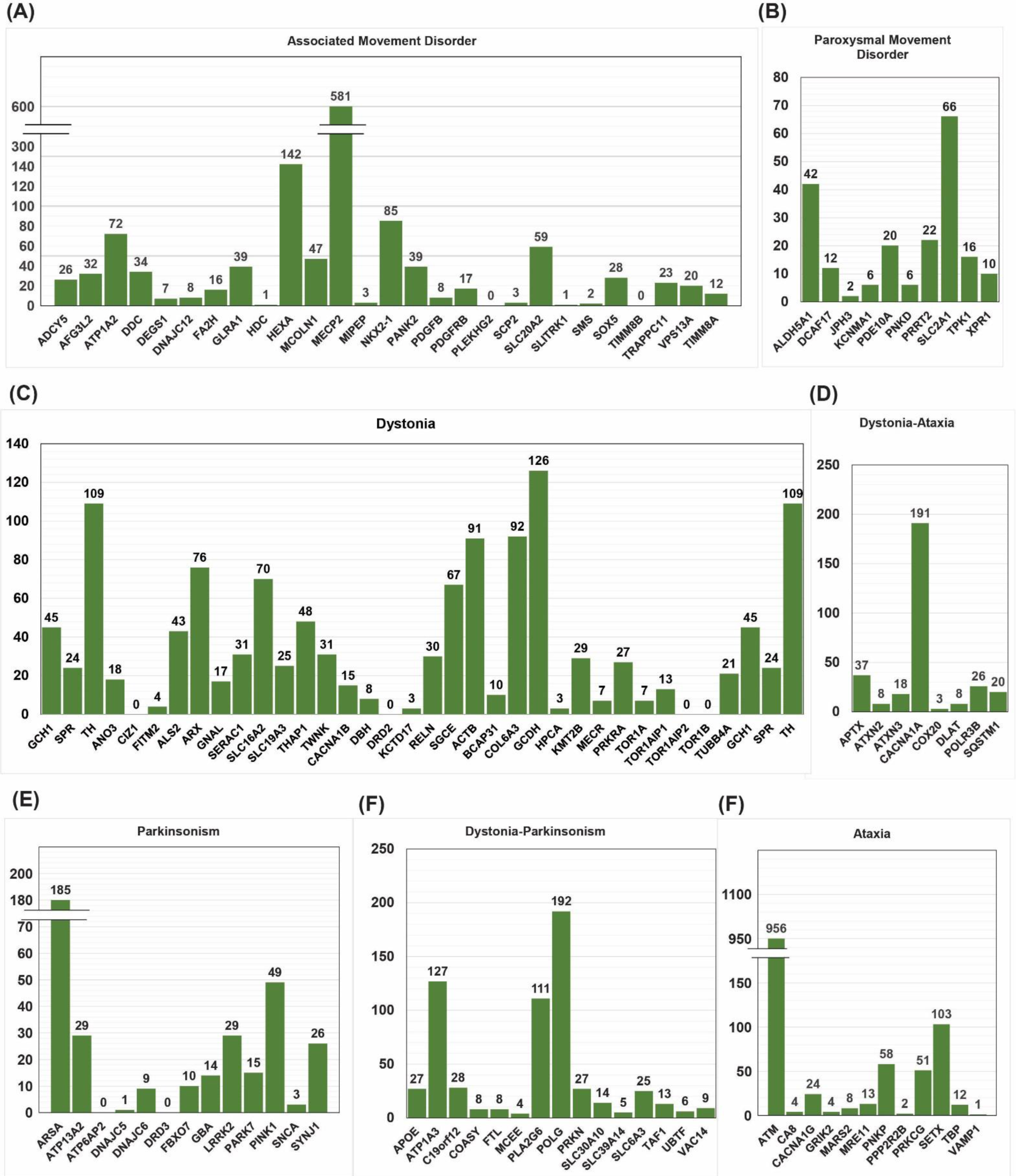
Number of pathogenic and likely pathogenic variants for gene clusters associated with movement disorder.

### Variant frequencies - Global populations overview

From a total of 28377 curated variants, 5118 variants were categorized as pathogenic and likely pathogenic. Of the total P/LP variants, 22 variants spanning 16 genes (ATP13A2, PINK1, PNKD, TRAPPC11, GLRA1, PRKN, DDC, TPK1, TPK2, ATM, HEXA, POLG, MCOLN1, GCDH, FBXO7 and ARSA) were found in the Indian population. Minor allele frequencies of these variants were compared between the Indian population and various other major global populations. **Supplementary Data 3** enlists the variant frequencies and Figure 3 gives a schematic overview of the distribution of these variants among populations.

**Figure 3:**
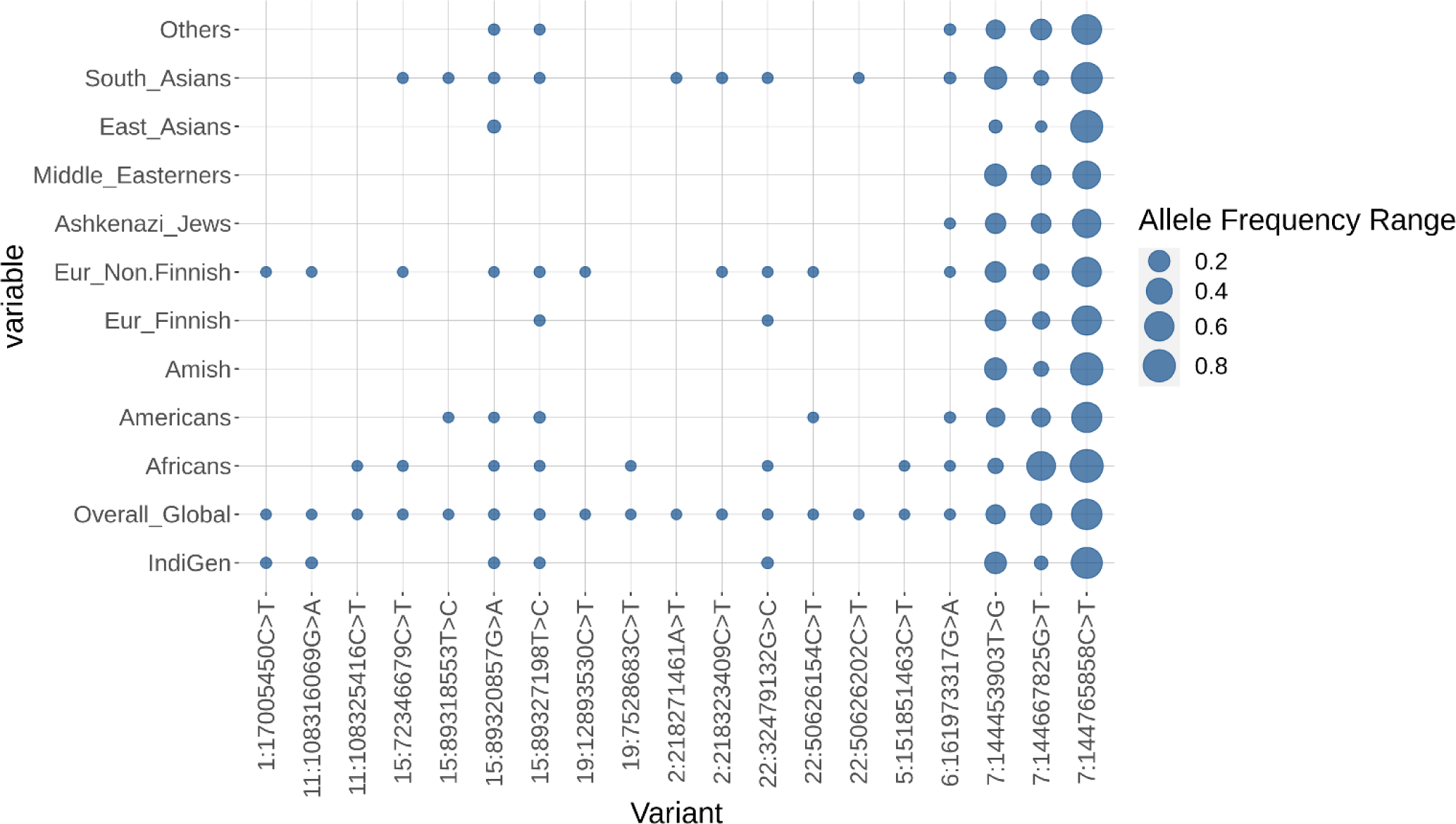
Comparison of pathogenic/likely pathogenic variants allelic frequencies between Indian Population (IndiGen dataset) and various other major global population datasets. The differences in the frequencies of variants which were filtered as significantly distinct among the global populations is shown in the bubble plot.

To the best of our knowledge, this is the first report for genome-scale analysis of variants associated with movement disorder in the Indian population.

### Database and web server

The DystoGen search engine is specifically designed to find relevant information about variants from a panel of 118 dystonia-associated genes that can be publicly accessed through URL https://clingen.igib.res.in/dystogen/. The search engine has been designed in a way that allows users to search for specific variant information either by querying the Variant, ACMG, AAChange, and dbSNP ID. Upon correct input, the result is displayed below the search bar containing basic information, including Gene, HGVS NM, AAChange, dbSNP, Mutation effect, Population, and ACMG status is shown in Figure 4. Upon clicking an individual variant from the results table, a new window opens up which contains pieces of information about the variant, that is displayed in four broad categories: ***Gene*** (contains information such as chromosome, chromosomal position, HGVS nomenclature, amino acid change, etc.), ***Mutation effect*** (contains information about the type of mutation, variant type, inheritance, ACMG classification, SIFT Score, SIFT prediction (deleterious/tolerant), Polyphen2 HVAR score, Polyphen2 HVAR prediction (deleterious/tolerant) and CADD Phred score. Apart from that it also contains information about allelic frequencies of the variants from three global population databases: 1000 Genome project (1000G_All), Exome Aggregation Consortium (ExAC_Freq), and ESP6500si_ALL.), ***Population*** in which variant was reported, and finally the ***Reference*** from where the variant was curated. The new window results page is displayed in **Supplementary Data 4**.

**Figure 4:**
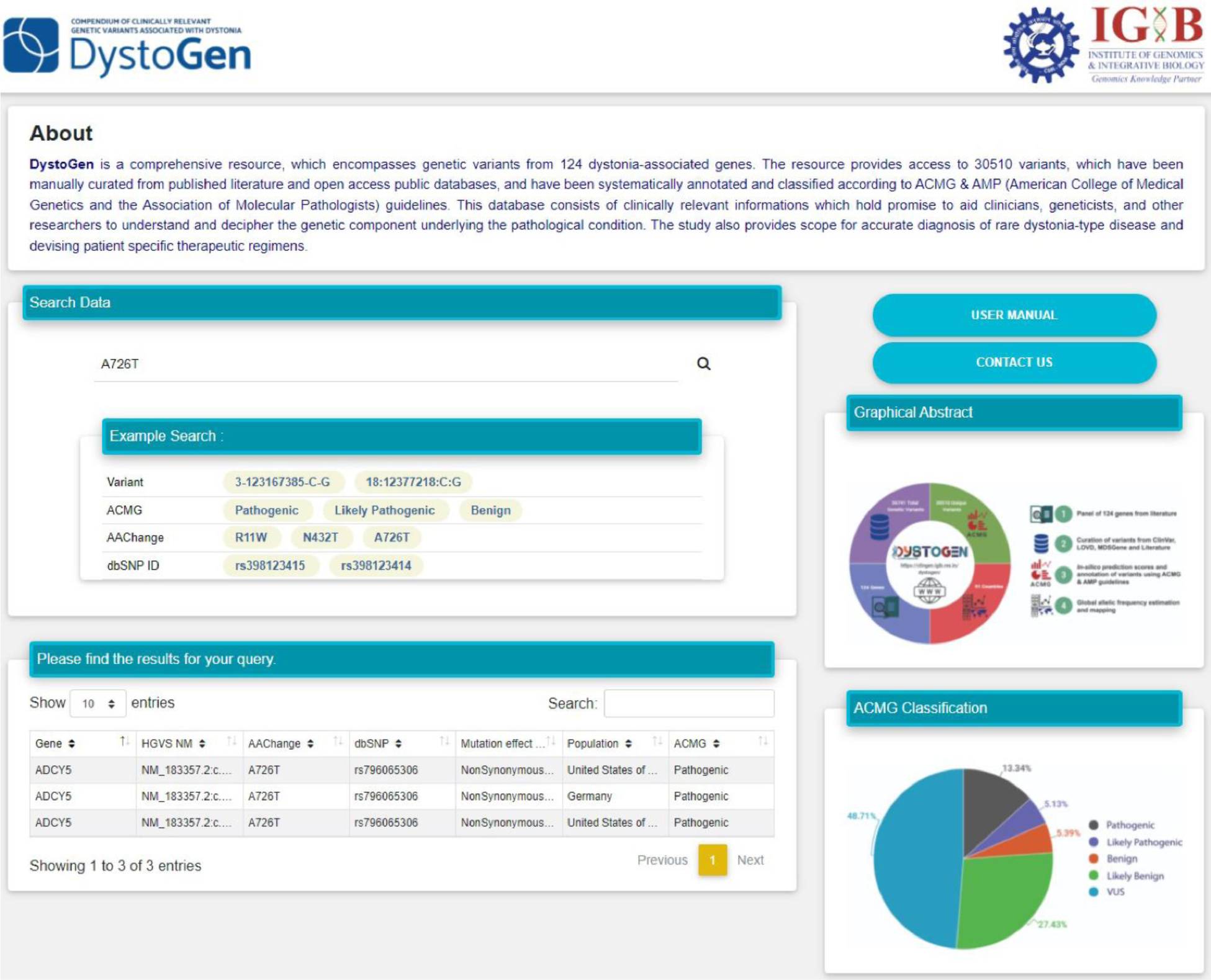
Result display against a search query.

## Discussion

Movement disorders comprise a diverse array of neurological conditions characterized by irregular and involuntary movements, or challenges in initiating, controlling, or coordinating movements. The diagnostic process for these disorders is intricate due to their varied presentations and complex genetic underpinnings. Relying solely on clinical assessment can be inadequate, particularly when symptoms are atypical or overlap with other conditions. This can lead to delays and inaccurate diagnoses, impacting patient care. Furthermore, traditional diagnostic techniques might miss underlying genetic factors when conventional genetic testing falls short.

Recent advances in Next Generation Sequencing (NGS) technologies have transformed diagnostics by enabling the identification of causative genetic variations. However, the sheer volume of genetic variants identified raises questions about their impact on health and their level of pathogenicity. The interpretation of genetic variants becomes pivotal for clinicians and researchers, informing drug development, therapeutics, and precise diagnosis.

An extensively curated and annotated database of genetic variants can play a central role in clinical diagnosis, enhancing disease management and aiding in understanding the link between genetic makeup and observable traits. In this context, evidence-based guidelines jointly established by ACMG & AMP offer a systematic approach to classify genetic variants, aiding clinicians in selecting patient-specific treatment strategies. These guidelines provide a structured framework to categorize variants based on their clinical significance, such as pathogenic, likely pathogenic, benign, or likely benign.

Building a database enriched with well-annotated information becomes a valuable resource for refining diagnostic accuracy, comprehending disease mechanisms, and advancing treatment methods. Such repositories expedite discovery and foster collaboration among diverse teams, propelling medical science forward. These resources not only hasten discoveries but also encourage interdisciplinary teamwork, driving progress in medical science, enriching our grasp of genetic diversity, and ultimately fostering innovative therapies to improve patient outcomes. The DystoGen resource (https://clingen.igib.res.in/dystogen/), serves as a unique platform for clinicians and researchers to swiftly analyze patients’ genetic variants, aiding disease diagnosis, treatment decisions, and family screening.

Although, ACMG guidelines have provided a standardized framework for variant interpretation, however, the limitations of these approaches, such as uncertain variants, variability in expressivity, involvement of modifier genes and unavailability of information regarding previously unreported variants renders difficulty to clinicians and researchers in confidently classifying a genetic variant. Apart from that, phenomenon such as balanced polymorphism, incomplete penetrance, founder effect and genetic drift further complicates the scenario. For instance, a well-studied LRRK2 mutation known as G2385R has gained extensive attention due to its recognized functional significance, making it a prevalent risk factor for sporadic Parkinson’s disease within the Chinese population (18). Notably, this variant has been identified in approximately 5% of the population, a classification that aligns with ACMG’s BS1 (Strong benign criteria), and has also received a benign classification from in silico prediction tools (BP4). Nevertheless, the biochemical assessment of LRRK2 G2385R in postmortem brain samples from Parkinson’s disease patients has revealed heightened kinase activity and synucleinopathy as distinctive pathological features (19). Despite its classification as benign according to ACMG guidelines, the presence of this mutation in a significant number of sporadic PD cases suggests a narrative that diverges from its classification.

While these genetic phenomena can complicate ACMG classification, it’s important to note that the ACMG guidelines are designed to incorporate a wide range of genetic scenarios. However, these challenges highlight the need for flexibility, expert judgment, comprehensive data integration in variant interpretation and development of robust platforms for analyzing the functional implication of such genetic changes. Geneticists and clinicians must carefully consider the nuances introduced by these phenomena, combine clinical and genetic information, and exercise caution in making variant classifications to ensure accurate and reliable clinical decision-making. Nonetheless, to the best of our knowledge, this is the largest database of genetic variants from different movement disorder associated genes, encompassing a wide range of populations across the world, that has been systematically classified according to the ACMG-AMP guidelines.

## Data availability

The data have been made available in the URL https://clingen.igib.res.in/dystogen/

## Acknowledgement

Authors acknowledge funding support from CSIR India through Grant RareGen, MLP2001, and a junior research fellowship to Bhaskar Jyoti Saikia from CSIR. Mukta Poojary acknowledges a junior research fellowship from DBT-BINC. We also acknowledge Mercy Rophina for data processing statistical analysis and global populations bubble plot. Funders had no role in the preparation of the manuscript or decision to publish.

## Author Information

These authors contributed equally: Bhaskar Jyoti Saikia and Utkarsh Gaharwar.

## Affiliation

**1. CSIR Institute of Genomics and Integrative Biology, Mall Road, New Delhi - 110007**

Bhaskar Jyoti Saikia, Utkarsh Gaharwar, Mukta Poojary, Aditi Mhaske, Sangita Paul, Mukesh Kumar, Srishti Sharma, Kavita Pandhare, Shivanshi Rastogi, Mercy Rophina, Vinod Scaria, Binukumar BK

**2. Academy of Scientific and Innovative Research (AcSIR), Ghaziabad, Uttar Pradesh 201002, India.**

Bhaskar Jyoti Saikia, Sangita Paul, Mukesh Kumar, Srishti Sharma, Mukta Poojary, Mercy Rophina, Vinod Scaria, Binukumar BK.

## Author contribution

Conceptualization: B.K, Data curation: B.J.S., U.G., A.M., S.P., S.R., M.K. Formal analysis: B.J.S., U.G., M.P., S.S. Funding acquisition: B.K, V.S., Investigation: B.K, V.S., B.J.S., U.G., A.M., S.P., S.R., M.K.. Methodology: V.S., B.K., B.J.S., U.G., M.P., K.P. Resources: V.S., B.K. Software: K.P. Supervision: V.S, B.K. Validation: V.S., B.K., B.J.S., U.G., A.M. Visualization: B.K., B.J.S., U.G., A.M., M.P., S.S. Writing—original draft: B.K., VS, B.J.S., Writing—review & editing: V.S., B.K., B.J.S., U.G., M.P., A.M., S.P., M.K., S.S., S.R.

## Supplementary information

***Supplementary Data 1:*** *List of genes selected for ACMG Classification along with associated movement disorder and its respective references*.

***Supplementary Data 2:*** *Individual gene clusters with pathogenic/likely pathogenic variants of each gene*.

***Supplementary Data 3:*** *List of pathogenic/likely pathogenic variants used for comparison of allelic frequencies between Indian Population (IndiGen dataset) and various other major global population datasets*.

**Supplementary Data 4:**
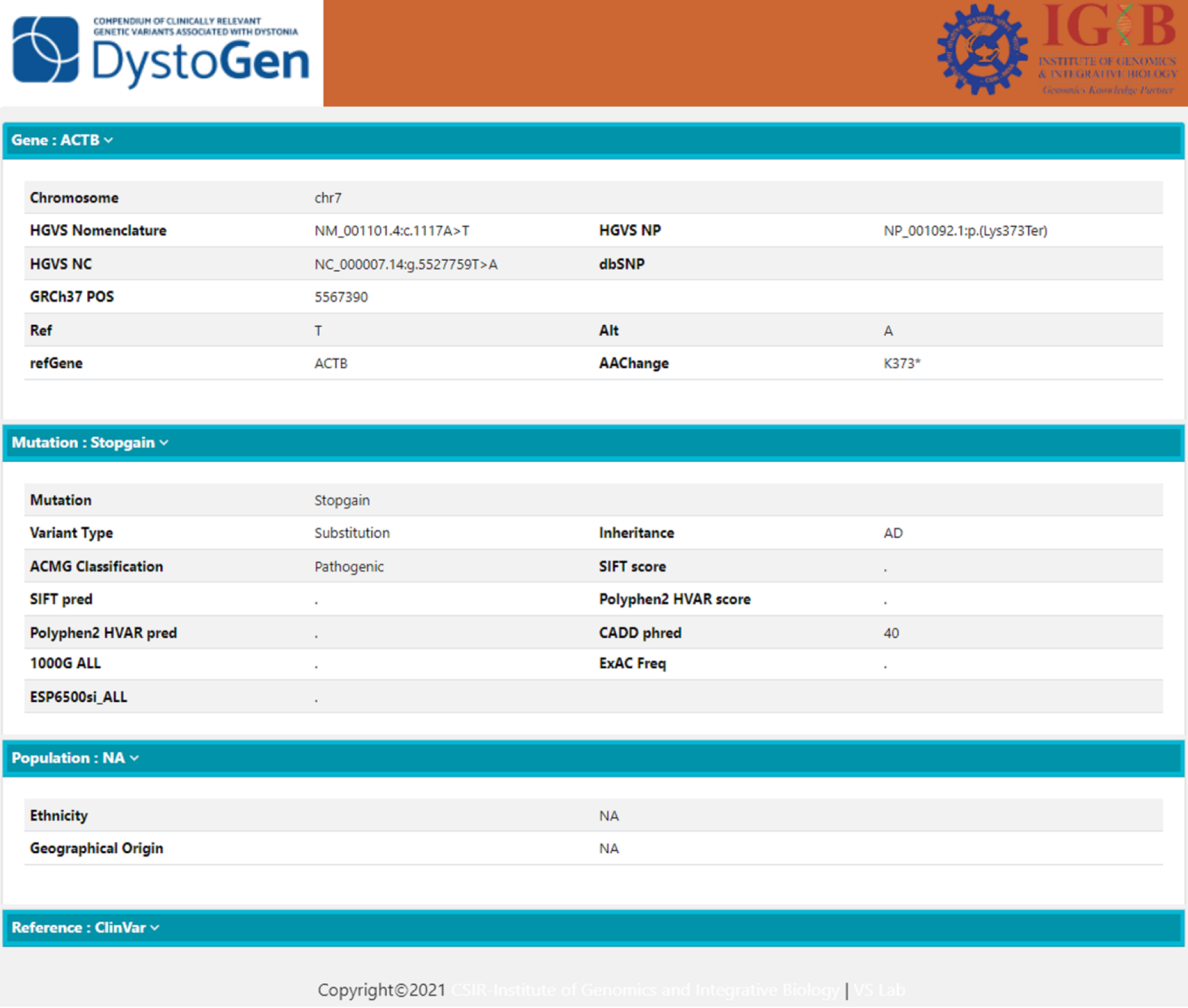
Result display against a search query.

