## Supplementary figures and images for "DystoGen Compendium: A comprehensive resource of ACMG annotated movement disorder associated genetic variants"

### Supplementary Data 4

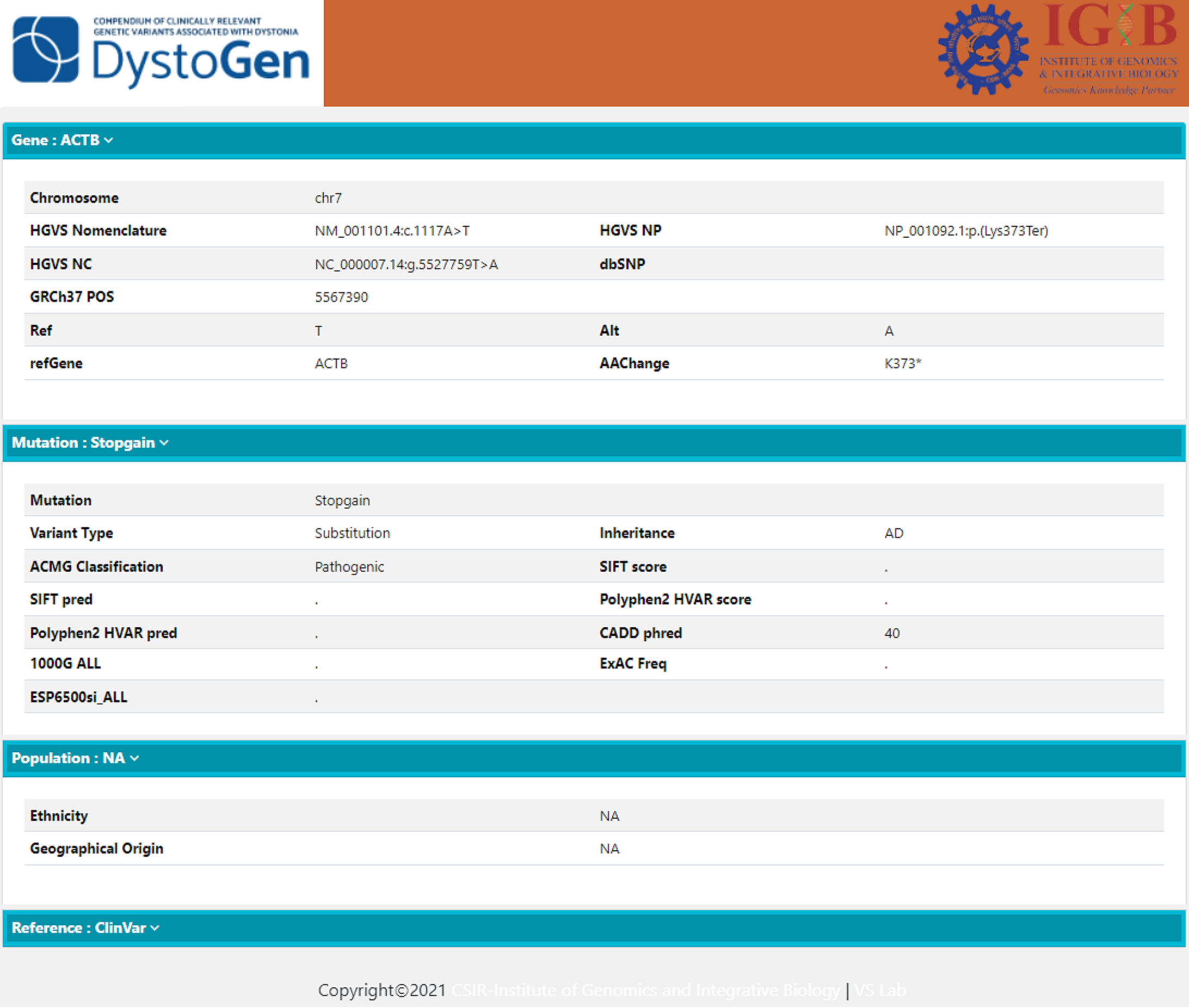
